# Epidemic Potential for Human Infection with Influenza A (H7N9) Virus in China through Web Search Behaviors: A Data-Driven Study

**DOI:** 10.1101/168112

**Authors:** Yue Teng, Dehua Bi, Xiaocan Guo, Dan Feng, Yigang Tong

**Author notes:** Correspondence: Yue Teng PhD, MD, State Key Laboratory of Pathogen and Biosecurity, Beijing Institute of Microbiology and Epidemiology, 20 Dong-Da Street, Fengtai District, Beijing 100071, China.; Yigang Tong Ph.D., State Key Laboratory of Pathogen and Biosecurity, Beijing Institute of Microbiology and Epidemiology, 20 Dong-Da Street, Fengtai District, Beijing 100071, China. These authors contributed equally to this work (Yue Teng, Dehua Bi).

## Abstract

Since the beginning of September 2016, a steep upsurge of the human cases of avian influenza A (H7N9) virus has been reported in China, which are alarming public concern for the pandemic potential of the H7N9 virus. In this study, we collected the data from H7N9 epidemics and H7N9-related Baidu Search Index (BSI) in China between September 2013 and June 2017. And we observed a strong correlation between the numbers of Influenza A (H7N9) cases and H7N9-related BSI in Guangdong province and Shanghai municipality (*p<0.001*). Autoregressive integrated moving average (ARIMA) models were constructed for the dynamic estimation of seasonal H7N9 outbreaks in 2016-2017 and the online search data acted as an external regressor with the historical H7N9 epidemic data in the forecasting model to improve the quality of predictions. Predictions by the models closely matched the actual numbers of reported cases during current H7N9 epidemic season. Especially, the estimated numbers of reported cases sharply increased to reach 49.88 (95% CI: 0-194.05) in Guangdong and 9.05 (95% CI: 0-37.43) in Shanghai from December 2016 to June 2017. Moreover, this accessible and flexible dynamic forecast model could be used in the monitoring of H7N9 virus to provide advanced warning of future emerging infection diseases.

**Author summary:** As the availability and popularity of the internet has greatly increased in recent years, an increasing number of cyber users, including patients and their family members, search online for health information on personal computers (PCs) and mobile phones (MPs) before seeking medical attention, making it possible to investigate the influenza prevalence by monitoring changes in frequencies of uses of particular search terms. In this study, we collected the data from H7N9 epidemics and H7N9-related Baidu Search Index (BSI) in China between September 2013 and June 2017. And then, we showed a strong correlation between the numbers of Influenza A (H7N9) cases and H7N9-related BSI in Guangdong province and Shanghai municipality (*p<0.001*). Furthermore, we reconstructed an improved dynamic forecasting method for outbreaks of H7N9 influenza using Autoregressive integrated moving average (ARIMA) models to predict future patterns of H7N9 transmission and the online search data acted as an external regressor with the historical H7N9 epidemic data in the forecasting model to improve the quality of predictions. Our results suggest that data from the Baidu search engine, combed with data from a traditional disease surveillance system, may be considered for early detection of H7N9 influenza outbreaks in mainland China.

## Introduction

The first case of avian influenza A(H7N9) virus in humans was reported in 2013[1, 2], and from then until July 2017, four previous seasonal outbreaks have been observed in China (the first season from approximately February to August 2013; the second season from September 2013 to August 2014; the third season from September 2013 to August 2014; the fourth season from September 2013 to August 2014; the fifth season from September 2016 to July 2017)[3-8]. There has been a significant increase in the numbers of reported human cases of H7N9 influenza at the end of 2016[7, 8]. Prevention and control of H7N9 influenza focuses primarily on traditional surveillance systems used for human infection with H7N9 influenza virus, which is based on passive or sentinel site monitoring in outpatient services and hospitals[9]. However, early and real-time surveillance of infectious disease prevalence, when followed by a rapid response, can reduce the effects of disease outbreaks[10-12].

The availability and popularity of the internet has greatly increased in recent years. At the same time, an increasing number of cyber users, including patients and their family members, search online for health information on personal computers (PCs) and mobile phones (MPs) before seeking medical attention, making it possible to investigate the health status of the population by tracking changes in frequencies of uses of particular search terms[13]. Surveillance of online behavior through monitoring queries in search engines is a potential web-based disease detection system that can improve monitoring. Google Trends (GT) has been shown to have the potential to go beyond early detection and forecast future influenza and Zika virus outbreaks[14, 15]. Several studies have used autoregressive integrated moving average (ARIMA) models for the forecasting of influenza prevalence from Google Flu Trends[16]. Baidu is the most popular search engine in China, making the Baidu Search Index (BSI) the most representative for analyzing online behavior and the awareness of cyber users in China[17]. The real-time nature of GT surveillance and the demonstrated strong correlation of GT data with infectious disease incidence suggest that GT may be a useful tool for early epidemic detection and prevention[18]. However, the forecasting capabilities of BSI for outbreaks of H7N9 influenza remain unknown. In this study, we examined H7N9-related BSI search terms temporally correlated with H7N9 influenza epidemics and developed an improved dynamic forecasting method for outbreaks of H7N9 influenza using ARIMA models to predict future patterns of H7N9 transmission.

## Materials and Methods

### Data collection and statistical analysis

The BSI (http://index.baidu.com) is an online system that tracks internet search volumes in China. BSI data were used to explore daily and weekly web behavior and cyber user awareness related to the H7N9 influenza outbreaks at city, province, and national levels. We collected the weekly BSI for search key word “H7N9” from personal computers (PCs) and mobile phones (MPs) in provincial level from 2 September 2013 to 19 June 2017 to cover the history of the H7N9 epidemic in mainland China (Table S1). The numbers of reported H7N9 cases per week at provincial level in mainland China from September 2013 to June 2017 were retrieved from the Food and Agriculture Organization (FAO;http://www.fao.org/AG/AGAINFO/programmes/en/empres/H7N9/wave_5/Situation_update_2017_06_28.html)and the Hong Kong Centre for Health Protection (CHP; http://www.chp.gov.hk/en/guideline1_year/441/332.html). Numbers of H7N9 confirmed cases per day with the illness onset date between September 2016 and June 2017 were obtained from WHO[7]; estimates of weekly incidence were converted from daily data (Table S1). To detect the BSI volumes associated with reported H7N9 cases, the Pearson product-moment correlation coefficient was used to assess linear correlation. All analyses were performed using Python v2·7 with the Scipy library.

### Reconstructed ARIMA model

For the time series analysis, autoregressive integrated moving average (ARIMA) models were fitted using R v2·14 software (R foundation for Statistical Computing, Vienna, Austria; http://www.r-project.org/). ARIMA (0, 1, 1) and ARIMA (2, 1, 1) forecasting models were developed for the Guangdong province and Shanghai municipality datasets, respectively (Table S1). We selected data from September 2013 to August 2015 as training set for model to learn and the learning model was tested with data from September 2015 to August 2016. To choose model parameters, the Box-Jenkins approach was applied to ARIMA (p, d, q) modeling of time series. This model-building process was designed to take advantage of associations in the sequentially lagged relationship that usually exists in periodically collected data[19, 20]. The parameter for the model included p, the order of auto-regression; d, the degree of difference; and q, the order of moving average. Several models were initially considered (Table S2); residual analysis and the Akaike Information Criterion (AIC) were used to compare the goodness-of-fit among the ARIMA models. The Ljung-Box test was used to measure the autocorrelation function (ACF) of the residuals (Table S2). Using this model selection procedure, we selected the ARIMA (0, 1, 1) model and the ARIMA (2, 1, 1) model to analysis the epidemic situations in the Guangdong province and the Shanghai municipality, respectively. To predict future values, the developed ARIMA models were fitted separately to the Guangdong province and Shanghai municipality datasets (Table S2) from 2 September 2013 to 4 September 2016 and used to forecast over a time span of 41 weeks (5 September 2016 to 19 June 2017).

## Results

### Correlation between reported human cases of H7N9 infection and cyber user awareness

Trends in the numbers of reported human H7N9 infections per province from 2 September 2013 to19 June 2017 and corresponding BSI data are shown in Figure 1 and 2. During the study period, three previous epidemics (the second, third and fourth H7N9 season) have been observed in China from September 2013 to August 2016: the one wave from October 2013 to August 2014 followed by the other outbreak from approximately September 2014 to July 2015 (Figure 1A, 2A and 2B). In late 2015 and early 2016, another wave of H7N9 infections has occurred (Figure 1A and 2C), and as of early 2017, the reported case counts began to increase again (Figure 1A and 2D), with 111 cases being reported from August 2016 onwards, and 100 cases being reported from in the first month of 2017 alone (Figure 2D). Since September 2016, not only has the fifth H7N9 outbreak started earlier than usual, but a steep increase in the number of humans infected with H7N9 virus has also been observed in Figure 1A and Figure 2D, causing domestic and international concern. According to the tendency of reported cases, similar trends were observed between numbers of human H7N9 cases and the BSI for H7N9 per month at provincial level, with search volumes peaking around January and February each year from 2014–2017 (Figures 1B and 2E-2H). Pearson product-moment correlation coefficients were calculated to examine temporal correlations between the H7N9 search queries and the reported numbers of H7N9 cases in Guangdong province (Figure 3A) and Shanghai municipality (Figure 3B). Analysis using the Pearson product-moment correlation coefficient suggested that H7N9 BSI from PCs and MPs were positively temporally correlated with the weekly numbers of reported cases in both Guangdong province (*p<0·001; R = 0·777;* Figure 3A) and Shanghai municipality (*p<0·001, R = 0·611;* Figure 3B), respectively.

**Figure 1.**
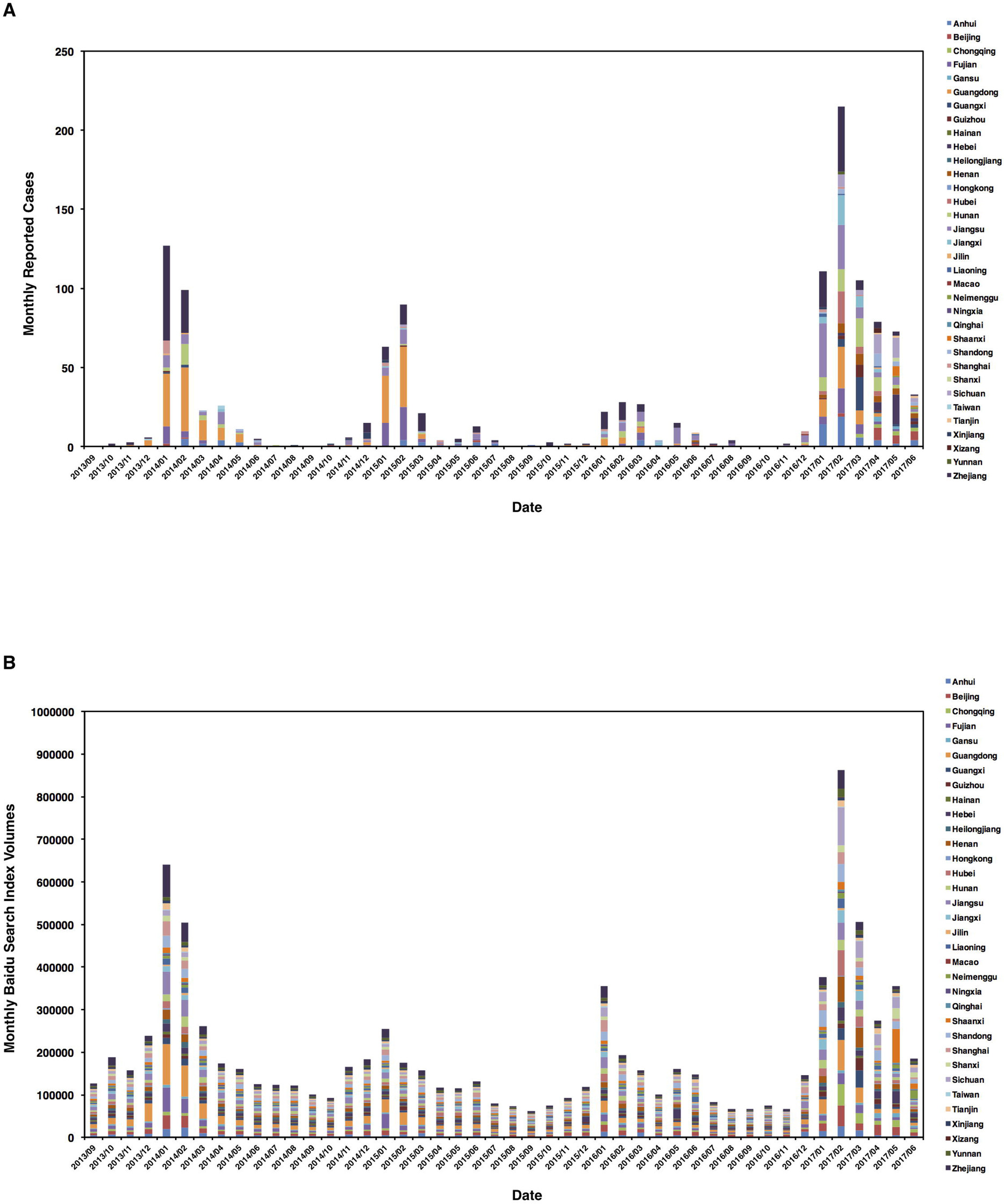
Time series plot of the reported cases per month of H7N9 influenza (A) and BSI volumes (B) in mainland China from September 2013 to June 2017.

**Figure 2.**
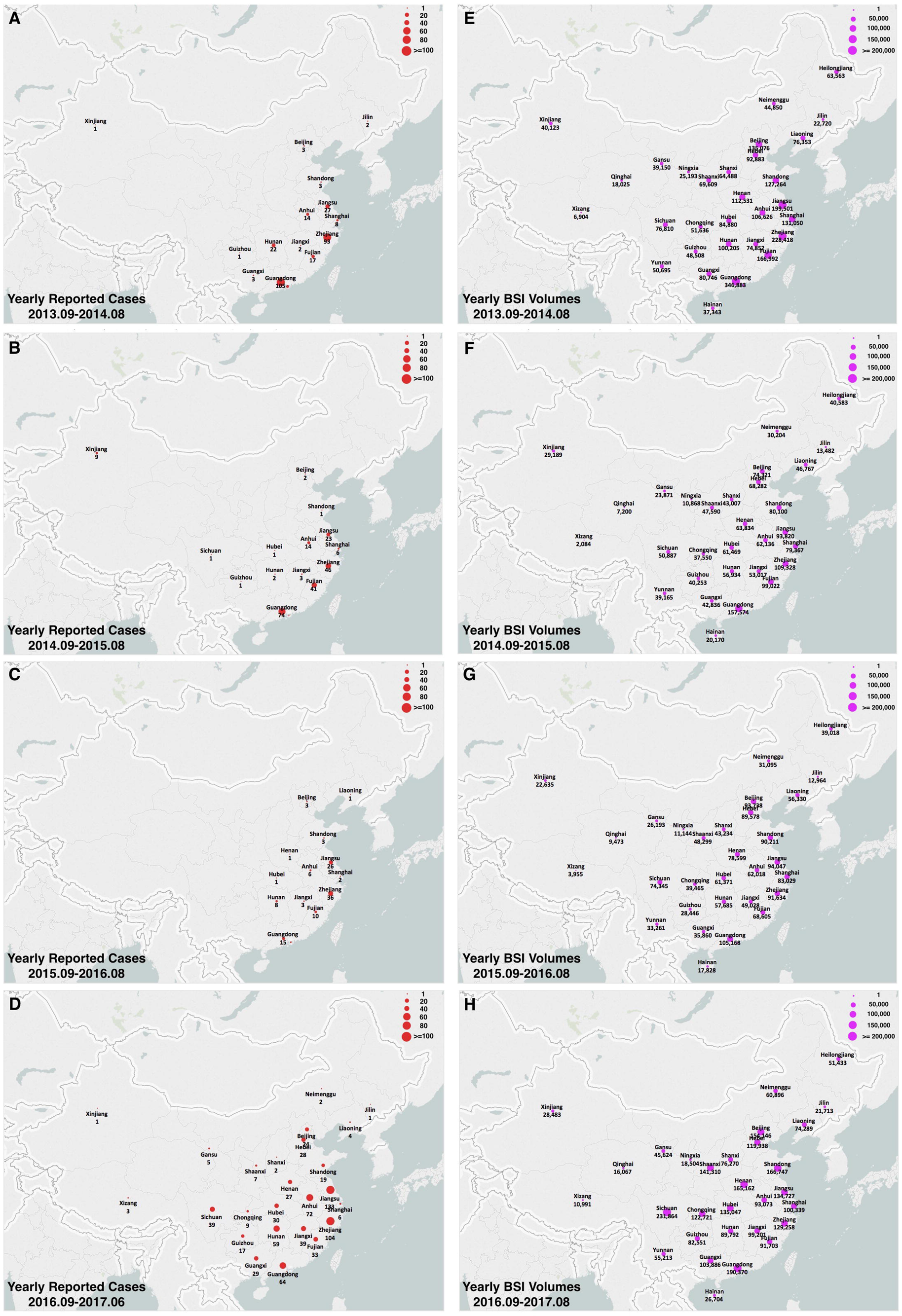
Spatial distribution of the reported cases per year of H7N9 influenza and BSI volumes in mainland China from September 2013 to June 2017. The darker red indicates a higher number of reported cases as well as a higher volume of BSI searches. The map was created in ArcGIS 9.2 software (ESRI Inc., Redlands, CA, USA).

**Figure 3.**
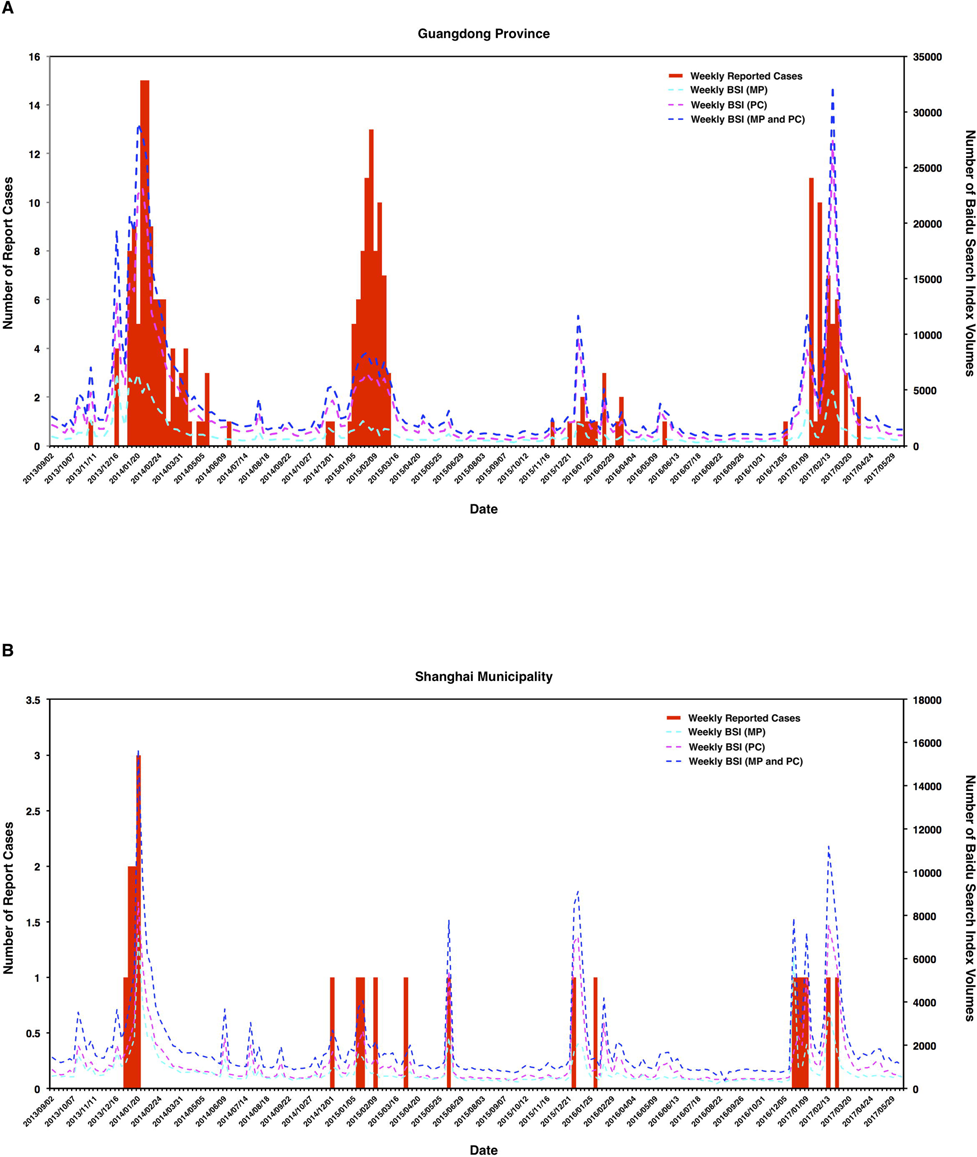
Distribution of reported cases per week of H7N9 influenza and BSI volumes in Guangdong province (A) and Shanghai municipality (B) from September 2013 to June 2017.

### Machine learning-based data-driven analysis

Based on the observed correlation between BSI search volumes and the numbers of reported cases, we divide the data into a learning set and a testing set. For both regions, data from September 2013 to August 2015 were used to train the model and data from September 2015 to August 2016 used to test it. Those models were fitted with the training datasets of Guangdong province and Shanghai municipality to observe whether the predictions were similar to the reported data in the testing set generated by the advanced ARIMA (0, 1, 1) model and ARIMA (2, 1, 1) model for the two regions in China, respectively. In both reconstructed models, the online search data were used as the external regressor to improve the predictions made from data on historic H7N9 epidemics. When both models were tested, with the training BSI data as a predictor, the 95% confidence interval (the blue regions in Figure 4) of the estimated number of cases included the number of actual reported cases in Guangdong province (Figure 4A) and Shanghai municipality (Figure 4B). In general, therefore, the predictions of both models were in accordance with the actual numbers of reported cases. These observations implied that the inclusion of BSI data could improve the prediction of the size of an outbreak of H7N9 influenza.

**Figure 4.**
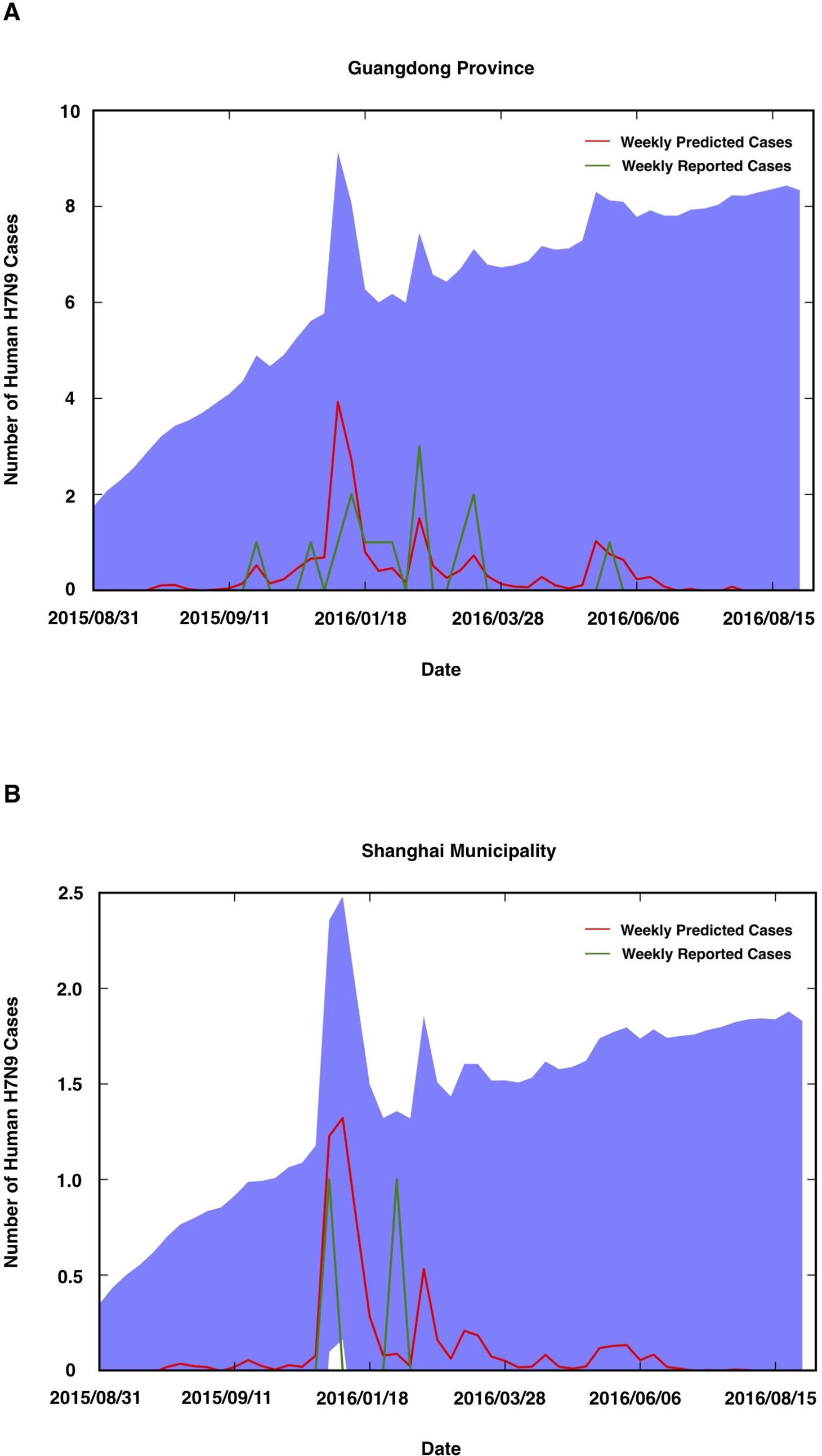
Numbers of reported cases of H7N9 influenza in the testing set compared with the simulation data generated by the advanced ARIMA (0, 1, 1) model and ARIMA (2, 1, 1) model for training set using BSI data as the external regressor for Guangdong province (A) and Shanghai municipality (B), respectively. The red and green solid lines represent the predicted and actual number of cases, respectively. The blue region represents the 95% confidence interval predicted by both models.

### Nowcasting the H7N9 outbreaks in the fifth epidemic during 2016 to 2017

Depending on the BSI data regarding H7N9-related online searches and epidemics of H7N9 influenza between 2 September 2013 and 4 September 2016, we used the reconstructed ARIMA (0, 1, 1) model and the ARIMA (2, 1, 1) model to forecast H7N9 cases in the first 41 weeks (from 5 September 2016 to 19 June 2017) of the 2016-17 H7N9 epidemics in the Guangdong province and the Shanghai municipality, respectively. The data regarding online searches were used as the external regressor to enhance the forecasting models and improve the quality of predictions based on data from historic H7N9 epidemics. The models estimated that 49.88 (95% CI 0–194.05; the blue regions in Figure 5A) and 9.05 (95% CI 0–37.43; the blue regions in Figure 5B) cases of human H7N9 influenza would be reported in Guangdong province (Figure 5A) and Shanghai municipality (Figure 5B), respectively, from 5 December 2016 to 19 June 2017. This matched closely with the actual numbers of cases reported (50 in Guangdong and 6 in Shanghai between 12 December 2016 and 19 June 2017). In addition, Figure 6 illustrated the seasonal geographical distribution of predicted human cases with H7N9 virus infection in Guangdong province and Shanghai municipal between September 2016 and June 2017. These forecasts suggest that the transmission of H7N9 influenza would be intense in these areas during December 2016 and February 2017, decrease after March 2017.

**Figure 5.**
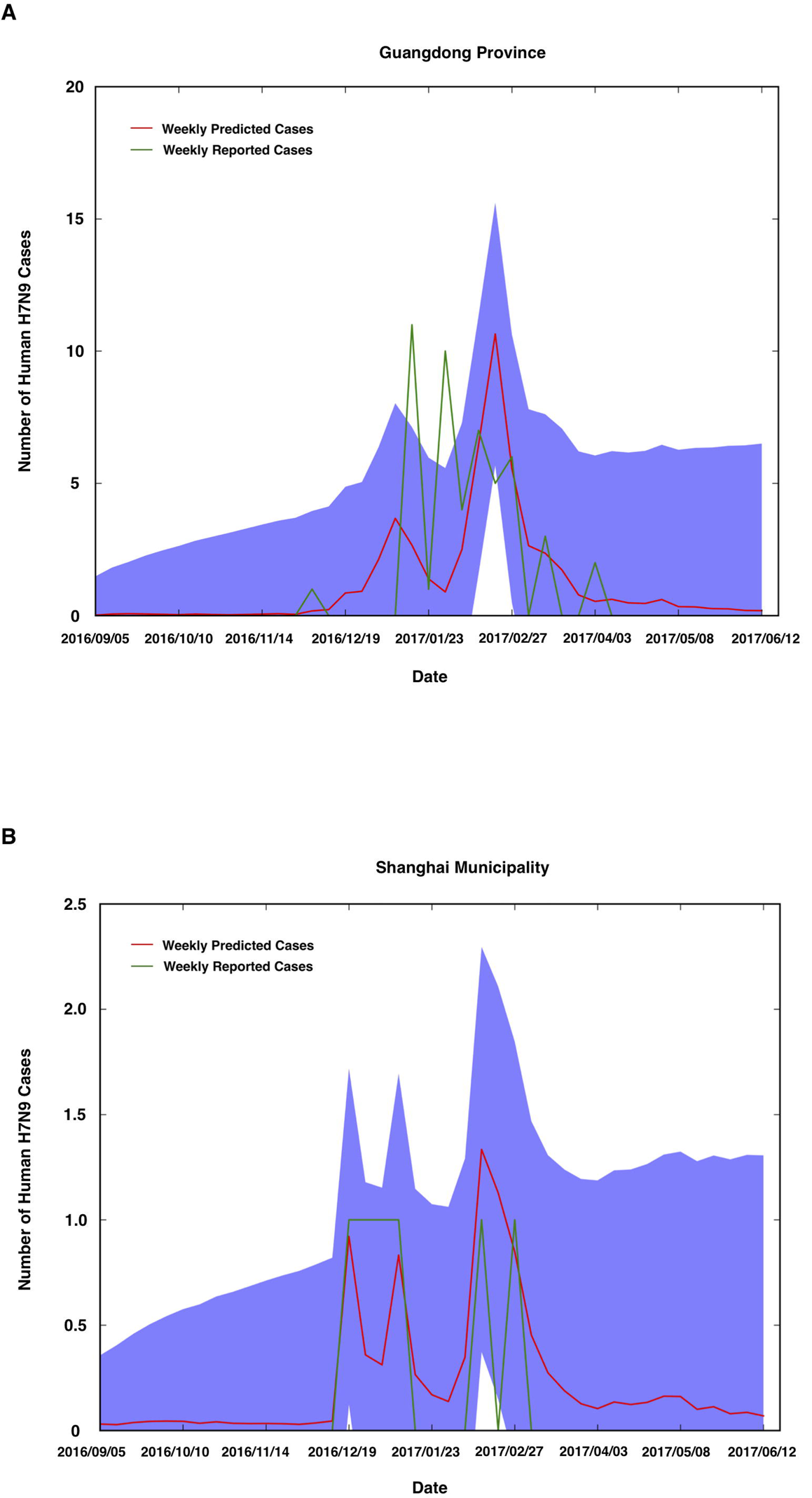
Forecasts of the numbers of cases of H7N9 influenza in Guangdong province (A) and Shanghai municipality (B) between October 2016 and June 2017 by the advanced ARIMA (0, 1, 1) and ARIMA (2, 1, 1) models, which were improved by aggregating historical logs with data of H7N9-related BSIs from the immediately preceding 10 months as an estimating predictor to estimate numbers of human cases of H7N9 infection. The red, green, and black solid lines represent the predicted, reported, and WHO-reported numbers of cases. The blue region represents the 95% confidence interval predicted by both models.

**Figure 6.**
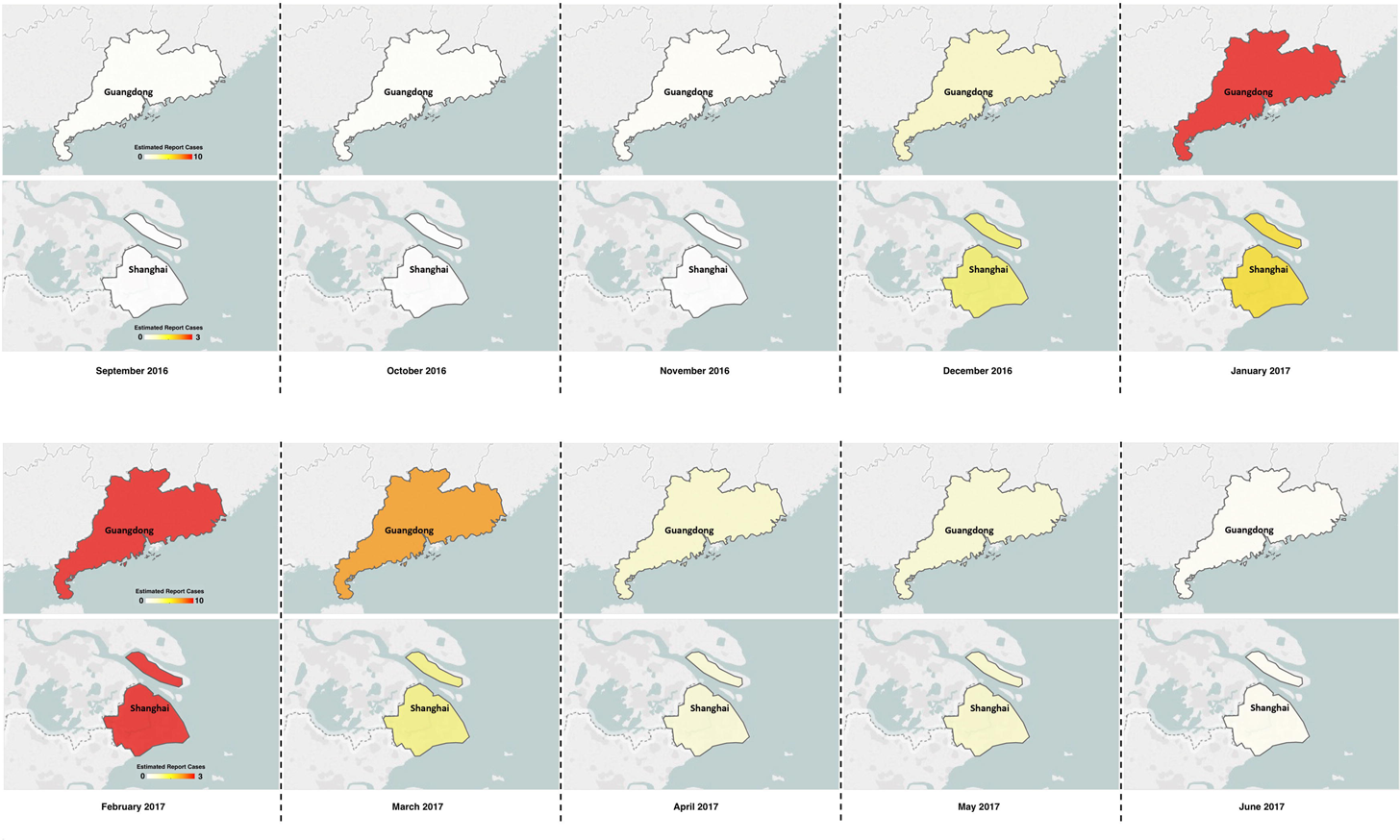
Geographical distribution of estimated numbers of reported cases per month of H7N9 influenza in Guangdong province (A) and Shanghai municipality (B) between September 2016 and June 2017. The map was created in ArcGIS 9.2 software (ESRI Inc., Redlands, CA, USA).

## Discussion

Outbreaks of human infection with avian influenza A (H7N9) virus in mainland China generally follow a seasonal trend, with four previous major outbreaks occurring to date[1-8]. However, the rapid increase in numbers of cases in the relatively short period since September 2016 has raised concerns about the pandemic potential of the H7N9 virus. As of 15 June 2017, there have been1533 cases of H7N9 influenza reported in China; WHO report that this includes at least 592 deaths[7]. Effective control of H7N9 virus outbreaks is contingent on high quality, timely surveillance data[2, 9, 21, 22]. Internet-based surveillance systems have therefore been increasingly explored as an innovative approach to improve the effectiveness of infectious disease prevention and control programs[15-17]. Data regarding cyber user awareness, in particular, are collected and processed in near real time from PCs and MPs; access to online search behavior produces monitoring data much faster than traditional systems.

This study combined novel and historical data on human cases of H7N9 influenza with sentinel surveillance data on cyber user awareness from BSI and showed a strong positive correlation between numbers of reported cases of H7N9 influenza and H7N9-related BSI (Figure 1-3). Based on the correlation data, the advanced ARIMA (0, 1, 1) model and ARIMA (2, 1, 1) model were improved by aggregating historical logs and estimates of H7N9-related search queries were used to forecast H7N9 epidemics in the Guangdong province and Shanghai municipality, respectively. The reconstructed ARIMA models performed well; predictions by the ARIMA models were similar to actual events during H7N9 outbreaks in Guangdong province and Shanghai municipality (Figure 4-6). This study also showed that data from the Baidu search engine, combed with data from a traditional disease surveillance system, may be considered for early detection of H7N9 influenza outbreaks in mainland China and may also be useful in providing dynamic timely information to public health agencies and near real-time indicators of the spread of infectious disease[23, 24]. However, further studies are needed that combine data from internet search queries and other potential factors (weather, live poultry, and contaminated environmental factors) to develop an early warning system[23, 24].

## Acknowledgments

We thank Dr. Wuchun Cao in State Key Laboratory of Pathogen and Biosecurity for discussions and our colleagues in the Beijing Institute of Microbiology and Epidemiology for help in technical assistance.

---

**Table S1.** Yearly Baidu Search Indexes and reported H7N9 cases at provincial level in mainland China September 2013 to June 2017.

**Table S2.** Comparison of tested models.

